# Cyanochelin uptake reveals an exclusively cyanobacterial class of AMIN-domain TonB-dependent transporters

**DOI:** 10.64898/2026.07.30.741596

**Authors:** Jan Mašek, Berness Peter Falcao, Lucie Kajan Grodecká, Vojtěch Hudzieczek, Tomáš Galica, Petra Urajová, Roman Sobotka, Lucie Kovářová, Roman Hobza, Pavel Hrouzek

**Author notes:** Correspondence to: Pavel Hrouzek, Roman Hobza.

## Abstract

Siderophore transport is central to microbial competition, because it determines access to iron, frequently a limiting nutrient. While siderophore-mediated iron uptake via TonB-dependent transporters (TBDTs) has been extensively studied in heterotrophic bacteria, little is known about the functionality and specificity of TBDTs in cyanobacteria. In the present study we functionally characterise the import system of cyanochelin B, a photolytic *β*-hydroxy aspartate siderophore produced by several filamentous cyanobacteria, including *Leptolyngbya* sp. NIES-3755.

We have identified a cyanochelin B putative transport cassete localized in the vicinity of the cyanochelin biosynthetic gene cluster in *Leptolyngbya* genome. By expressing the import genes heterologously in a model unicellular cyanobacterium *Synechocystis* sp. PCC 6803, we established that the transport cassette reconstitutes cyanochelin B-dependent growth, consistent with cyanochelin-mediated iron acquisition. Systematic gene dissection showed that the TBDT (CctA) and the substrate-binding protein (CctB), responsible for binding the siderophore in the periplasm, are alone sufficient for cyanochelin import into *Synechocystis* cells, with the permeases, ATPase and a cassette-associated ferredoxin supplied *in trans* by the host. CctA carries an N-terminal AMIN domain, a fusion found only in cyanobacterial TBDTs. The cassette accepts the structurally similar cyanochelin A but not cyanochelin C, enterobactin or pyoverdine, indicating limited promiscuity. The phylogenetic placement of cyanochelin receptors within a broader clade containing citrate-hydroxamate-type siderophore receptors suggests an evolutionary link between transport systems for chemically distinct cyanobacterial siderophores. Our study reports the first functional heterologous expression of a cyanobacterial TonB-dependent transporter and establishes *Synechocystis* as a promising platform for cyanobacterial xenosiderophore-uptake studies.

## Introduction

Iron is an essential micronutrient for cyanobacteria. It serves as a cofactor in various enzymes, notably in photosynthetic electron transfer and N_2_ fixation, which increases cellular demand as much as tenfold compared to heterotrophs.^1,2^ Despite being the fourth most abundant element in the Earth’s crust, most iron is present in the oxidised ferric (Fe^3+^) form that is generally insoluble and poorly biologically available. As a result, iron is a growth-limiting element in many ecological niches.^1,3^

Cyanobacteria possess several mechanisms for iron uptake. Ferric or ferrous iron can be transported into periplasmic space through porins in the outer membrane and subsequently into the cytoplasm via ABC transporters (e.g. FutABC) or iron transporters such as FeoB localized at the cytoplasmic membrane.^4–6^ Moreover, *Synechocystis* sp. PCC 6803 was also shown to be capable of reducing inorganic Fe(III) via a respiratory terminal oxidase in the periplasm.^6,7^

Like other bacteria, cyanobacteria employ siderophores for iron scavenging when iron starvation becames severe. These low-molecular weight compounds typically features strong iron-binding capabilities, which rely mostly on catechol, hydroxamate or α-hydroxycarboxylate moieties.^8,9^ Some of the best studied siderophores are enterobactin, pyoverdines or desferrioxamines. Cyanobacterial siderophores include schizokinen,^10^ anachelin,^11^ synechobactins,^12,13^ leptochelins^14^ and cyanochelins.^15,16^ Diverse structurally complex bacterial siderophores are often synthesised via nonribosomal peptide synthetases, sometimes merged with polyketide synthases.^8^

In microbial communities, siderophores play a pivotal role in microbial competition for iron. Along with siderophore production, successful transport of the ferri-siderophore complexes into the cell is an important aspect in microbial iron competition as the producer finally recoups the energetic cost of siderophore synthesis.

The most common pathway in Gram-negative bacteria comprises a TonB-dependent transporter (TBDT) at the outer membrane and an ABC transporter at the plasma membrane (see Figure 1A).^17,18^ The ferri-siderophore complex is imported in an energy-dependent manner from the outer environment into the periplasm by the TBDT, a 22-stranded β-barrel with a globular plug domain inside of the barrel. The energy from the plasma membrane’s proton motive force is provided by the interaction of a TBDT with the inner membrane protein TonB in complex with ExbB and ExbD. In the periplasm, the siderophore-iron complex is bound by a substrate-binding protein (SBP) and directed to an ABC type transporter that translocates it into the cytoplasm. The release of iron occurs mostly through siderophore hydrolysis, iron reduction with subsequent siderophore dissociation, or photolysis of the siderophore.^16,19^

**Figure 1.**
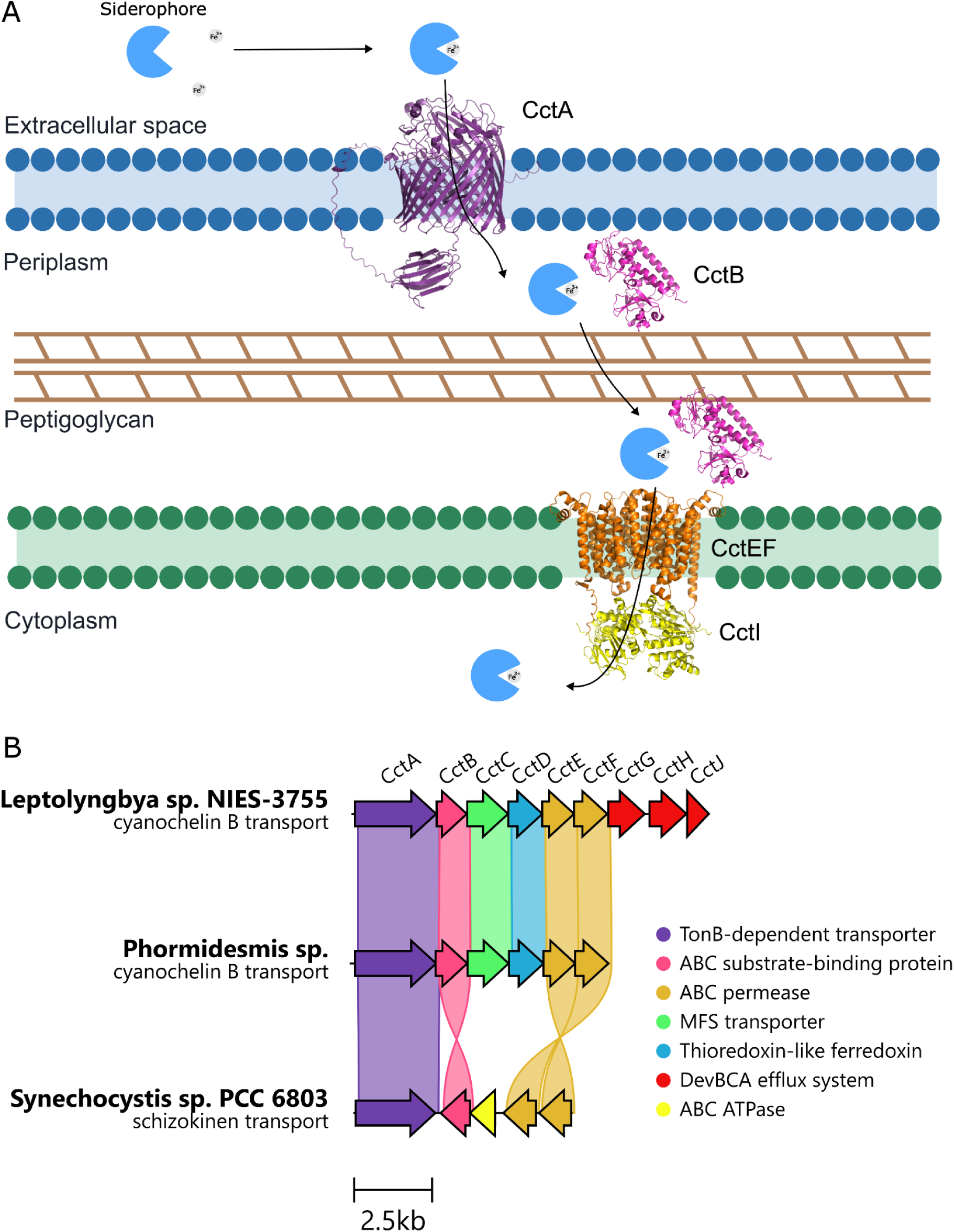
Cyanochelin transport machinery and transporter cassete organisation. **A**, **Suggested cyanochelin B import machinery.** The ferri-siderophore complex is translocated through outer membrane via the TBDT CctA. In the periplasm, the complex is bound by SBP CctB and directed to ABC transporter CctEFI. The CctI protein is encoded by a chromosomal gene. **B**, **Organisation of transport genes of cyanochelin B and schizokinen.** Upper two shown clusters are located immediately upstream of cyanochelin B BGC and share high level of protein identity. *Leptolyngbya* sp. NIES-3755 cassette contains additional DevBCA efflux system compared to *Phormidesmis* sp. Third cluster is schizokinen transport cassette FecABCDE in expression strain *Synechocystis* sp. PCC 6803 Nixon, having all ABC transporter components and lacking MFS exporter.

Besides siderophore producers, which express both biosynthetic and transport genes, there are also cheaters which express only the siderophore uptake machinery, without having to invest in siderophore production. Thus, the specificity of the import pathway, especially TBDT receptor, is crucial for the producer to benefit from the whole process. Microbial interactions concerning siderophores are well described in the review by Kramer *et al*.^9^

In recent years our research team discovered and characterised a novel group of cyanobacterial siderophores coined cyanochelins.^15,16,20^ Their defining structural feature is the presence of two *β*-hydroxy aspartate moieties involved in iron chelation. We experimentally determined structures of cyanochelins A, B and C and found that cyanochelins are probably widespread siderophores across the cyanobacterial phylum. Co-cultivation of a cyanochelin B producer with non-producing *Synechocystis* revealed that the benefit alternates between private monopolisation by the producer and community-wide benefit through photolytic cleavage of the siderophore.^16^ In the present study, we set out to experimentally determine properties of the cyanochelin B transport system. We show that a cassette of several genes immediately upstream of the cyanochelin biosynthetic gene cluster (BGC) is responsible for its transport. We determine its minimal functional components, its substrates and reveal an evolutionary relationship between cyanochelin and hydroxamate siderophore receptors.

## Results

### A self-contained cassette encodes the cyanochelin B transport cycle

A cluster of nine cyanochelin transport genes (*cctA-H, cctJ*) is situated on the complementary strand immediately upstream of the cyanochelin B BGC in *Leptolyngbya* sp. NIES-3755. The whole BGC and the transport cassette genes are localised on a ∼100 kbp plasmid. The manual inspection combined with prediction using antiSMASH identified them to be four import genes (*cctA*,*B*,*E* and *F*), four export genes (*cctC*,*G*,*H* and *J*), and one gene of unknown function (see Figure 1B + Table 1).

**Table 1.** Genes in the transport cassette of cyanochelin B.

| Gene | ID | Proposed annotation | Homolog | Homolog id | Homolog organism | Protein identity (%) |
| --- | --- | --- | --- | --- | --- | --- |
| cctA | BAU16017 | TonB-dependent transporter |  |  |  |  |
| cctB | BAU16018 | periplasmic unit of ABC transporter | fecB1 | slr1319 | Synechocystis sp. PCC 6803 | 34,7 |
| cctC | BAU16019 | Major Facilitator Superfamily protein | entS | P24077 | Escherichia coli K-12 | 22,4 |
| cctD | BAU16020 | thioredoxin-like sucrose ferredoxin |  |  |  |  |
| cctE | BAU16021 | permease unit of ABC transporter | fecC | slr1316 | Synechocystis sp. PCC 6803 | 55,3 |
| cctF | BAU16022 | permease unit of ABC transporter | fecD | slr1317 | Synechocystis sp. PCC 6803 | 58,4 |
| cctG | BAU16023 | HlyD family efflux transporter periplasmic adaptor subunit | DevB | all2652 | Anabaena sp. PCC 7120 | 60,7 |
| cctH | BAU16024 | FtsX-like permease family | DevC | all2651 | Anabaena sp. PCC 7120 | 62,2 |
| cctJ | BAU16025 | ATP-binding cassette domain-containing protein CDS | DevA | alr3649 | Anabaena sp. PCC 7120 | 69,2 |

The whole cassette starts with the first import protein CctA – a TonB-dependent transporter (BAU16017). The predicted structure (see Figure 2B) included two typical domains – a β-barrel and a plug domain situated in the barrel lumen, as well as an AMIN domain at the N-terminus of the protein between the signal sequence and the plug domain. The TBDT CctA is followed by the substrate-binding protein CctB and permease units of ABC transporter CctE and CctF. The 5’ end of *cctE* overlaps with the upstream *cctD* gene, and its 3’ end with *cctF*. The CctEF permease complex shows homology to *Synechocystis* FecCD permeases involved in schizokinen uptake^21^ (protein sequence identities 55.3 % and 58.3 % for CctE and CctF, respectively). Notably, the ATPase unit of the ABC transporter is absent. We found a homologous protein CctI (BAU12306.1; 63.2 % identity with *Synechocystis* FecE ATPase) on the *Leptolyngbya* chromosome, separated from plasmid-borne CctEF.

**Figure 2.**
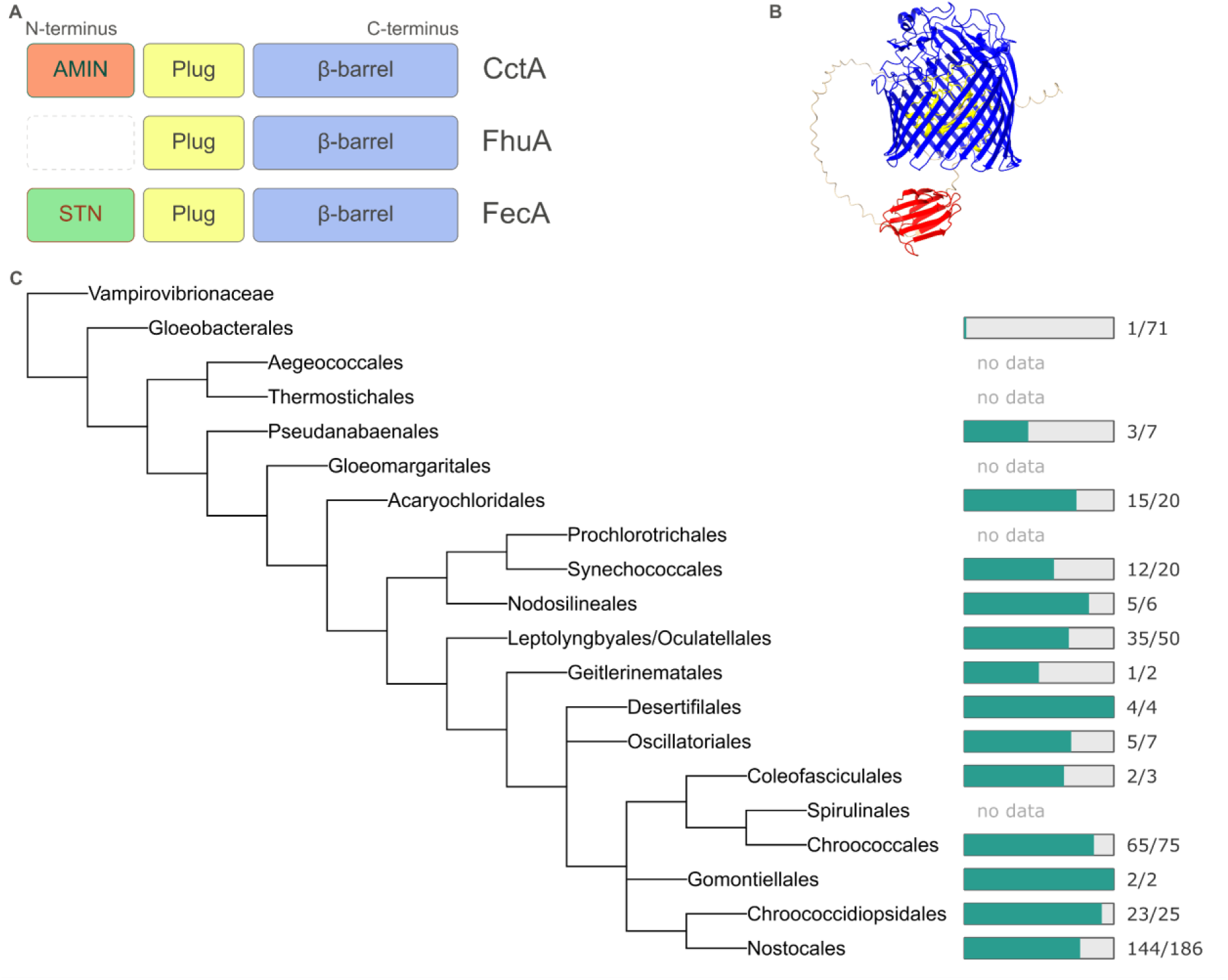
Architecture and distribution of AMIN domain in cyanobacteria. **A**, Domain architecture of TBDTs - *Leptolyngbya* NIES-3755 cyanochelin B receptor CctA, *E. coli* ferrichrome receptor FhuA and *E. coli* dicitrate receptor FecA. **B**, Prediction of CctA structure by AlphaFold Server.^29^ The AMIN domain is coloured red, the plug domain yellow and barrel domain dark blue. **C**, Distribution of AMIN domain through cyanobacteria. As dataset all complete cyanobacterial genomes in Refseq were used. The numbers show ratio of AMIN-TBDTs to all TBDTs from the taxa. Cladogram adapted from Strunecký *et al.*, 2023.^30^

There are two putative export systems in the gene cluster. CctC is a Major Facilitator Superfamily (MFS) protein with typical 12 α-helices topology.^22^ The last three proteins of the cassette, CctG, CctH and CctJ are homologues of *Anabaena* sp. PCC 7120 DevBCA efflux pump.^23^ As Falcao *et al.*^16^ pointed out, the cyanochelin B producer *Phormidesmis* sp. encodes full cyanochelin B BGC accompained with a cassete of only six transport genes where the second efflux system CctGHJ is absent (Figure 1). This suggests that MFS transporter might be sufficient for export of the siderophore into the periplasm.

The CctD (BAU16020) was not classified as a transport protein by antiSMASH. It is predicted to be part of the thioredoxin-like ferredoxin protein family. According to the InterPro database (IPR009737), this protein family is distributed across bacteria, fungi and plants and some members show sucrolytic activity. We noticed this class of protein to be present in other siderophore transport clusters throughout the cyanobacterial phylum, seen during manual inspections as well as in a CluSeek search (see Figure S1).^24^ Still, we lack sufficient evidence to assess its putative role, for example in iron reduction.

### AMIN-TBDT fusion is unique to cyanobacteria

The AMIN domain seemed to be an unusual feature of a TBDT receptor. It was firstly described in a bioinformatic paper in 2008, in which the authors found it to be widely associated with cell envelope proteins. The CctA’s architecture (AMIN-plug-barrel) was mentioned as one of many, without further investigation.^25^ In follow-up research it was proven to bind peptidoglycan and localise N-acetylmuramyl-L-alanine amidase AmiC in *E. coli*.^26^

Some TBDTs feature an additional domain at the N-terminus (see Figure 2A), most notable being the signal domain (STN) in ferric citrate transporter FecA in *E. coli* or ferric pyoverdine transporter FpvA in *Pseudomonas aeruginosa*. The STN regulates transcription of the transport gene operon through sigma factor activation/repression cascade.^27,28^ While the STN domain is well described in TBDTs, the AMIN domain is not present in typical transporters from *E. coli* or *P. aeruginosa*. To determine the prevalence of such a domain architecture, we used HMM profiles to scan all complete bacterial *in silico* proteomes in RefSeq for the presence of an AMIN domain in a TBDT. Out of 251,691 entries, only 395 contained the domain. Closer inspection revealed that all of them are of cyanobacterial origin. The AMIN-TBDT fusion therefore seems to be a unique cyanobacterial invention. We further investigated this finding by clustering the sequences at 90 % identity and examining the distribution of the AMIN-TBDT fusion through the phylum (Figure 2C). We discovered that 66 % (319/484) of the cyanobacterial TBDTs do have the domain. The domain is spread through most cyanobacterial taxa, even though it is worth noting that some orders had only a few members in the dataset. Notably, an exception was found in early-branching order *Gloeobacterales*: only 1 out of 71 TBDTs contains an AMIN domain. We inspected the TBDT and found it to be situated in a genomic island of other putative siderophore transporters, suggesting acquisition by horizontal gene transfer.

### Cloning and heterologous expression of the transport cassette

To determine experimentally whether the cassette mediates cyanochelin B uptake, we expressed it heterologously in *Synechocystis* sp. PCC 6803 (hereafter *Synechocystis*). *Synechocystis* is a well-established model, lacks siderophore production of its own and harbours only a few TBDTs.^31,32^

Initially, we attempted expressing the TBDT receptor CctA alone in the wild-type strain (WT); the resulting strain did not show decisive cyanochelin B import (data not shown). Our *in silico* analysis described above indicated that four of the genes (*cctA*, *cctB*, *cctE* and *cctF*) encode TBDT and ABC transporter components. We therefore constructed a vector carrying all four import genes together with *cctD*, whose function we could not assign but which was present in related transport cassettes. The ABC transporter ATPase, also required for import, is not colocalised with the other transport genes on the plasmid. We therefore expected that in the native cyanochelin producer, it is supplied by the chromosomal gene. Accordingly, we reasoned that the missing subunit could be complemented by a *Synechocystis* FecE protein and decided to test only the genes in the immediate vicinity of the cyanochelin BGC. As a surrogate host, we turned to the *Synechocystis ΔfutC* knock-out mutant.^5^ The *futC* encodes the ATP-binding unit of an ABC-type transporter responsible for Fe^3+^ uptake under iron starvation. We observed that *Synechocystis* WT is able to proliferate well even after long-term iron starvation, which partly hampered the functional assay relying on growth measurement. As the *ΔfutC* knock-out is substantially more sensitive to iron starvation, we selected it for the assay to provide a clean background. All five putative import genes were cloned into a pPD-FLAG based plasmid^33^ for constitutive expression and transformed into *Synechocystis*.

The resulting strain (CctABDEF) was cultivated in a 96-well plate exposed to cross-gradients of cyanochelin B and ferric iron FeCl_3_, and optical density at 750 nm was measured (see Figure 3A). The growth of the control *ΔfutC* strain was entirely abolished under iron starvation at all cyanochelin concentrations. Our strain CctABDEF proliferated poorly under iron limitation with no cyanochelin addition. Adding cyanochelin B resulted in prominent cell growth of CctABDEF compared to the *ΔfutC* control. This clearly indicates that the import cassette is able to compensate for iron starvation via the cyanochelin import into *Synechocystis* cells.

**Figure 3.**
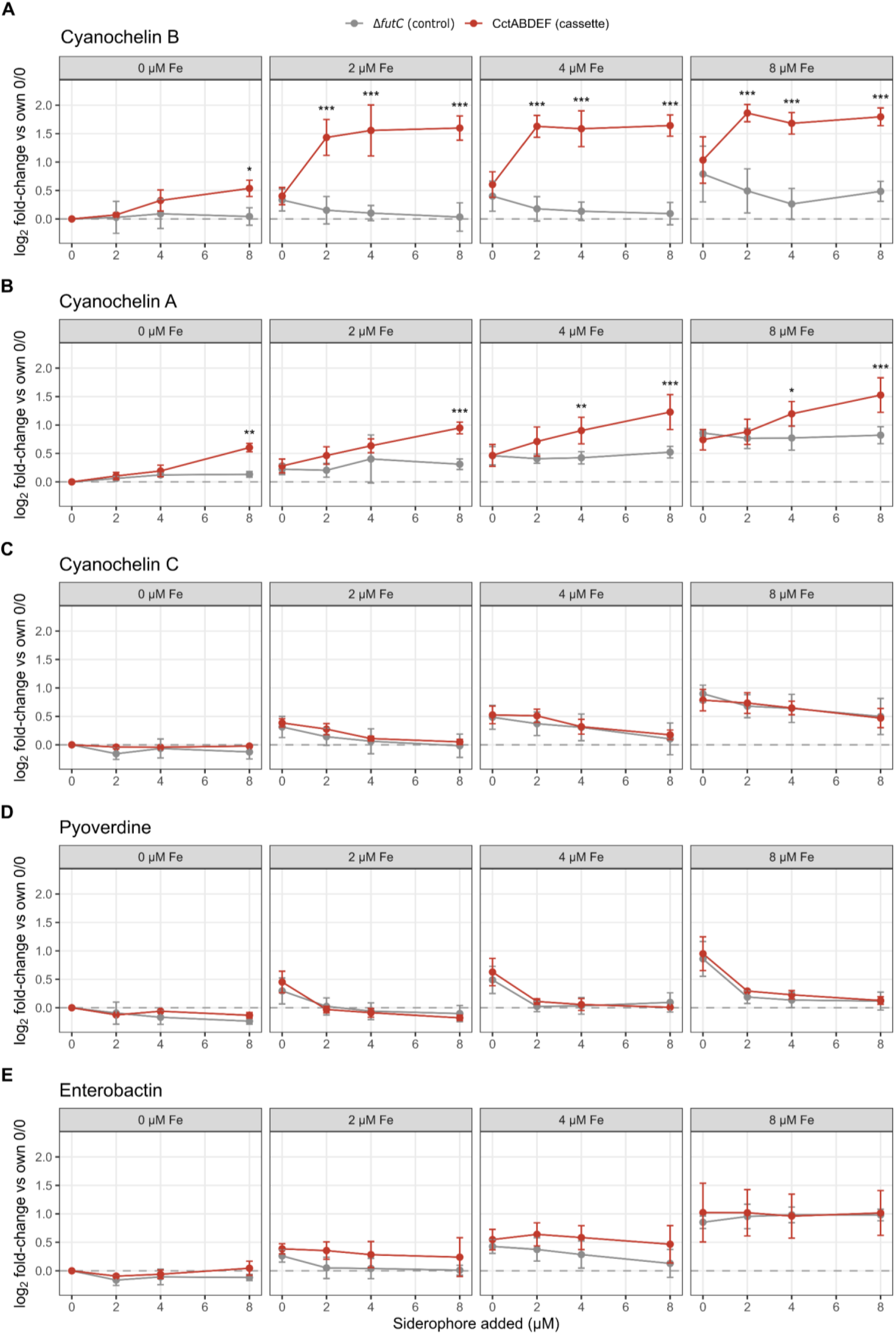
Substrate specificity of the cyanochelin B transport cassette across various (xeno)siderophores. For each siderophore, growth (OD₇ ₅ ₀ ) is shown as log₂ fold-change relative to each strain’s own iron- and siderophore-free (0/0) condition, computed within each biological replicate (n = 3); points are means, error bars ± SD, and the dashed line marks no induction. Panels A–E correspond to cyanochelin B, cyanochelin A, cyanochelin C, enterobactin and pyoverdine; within each panel, sub-panels are iron concentrations (0–8 µM), the x-axis is added siderophore (0–8 µM). Statistical significance reflects the difference in induction between the cassette strain and the control at each iron × siderophore combination, obtained from a linear model (log₂ fold-change) fitted per siderophore, with the cassette-minus-control contrast extracted by estimated marginal means and Benjamini–Hochberg correction across the fifteen cells of each siderophore (*P < 0.05, **P < 0.01, ***P < 0.001). Control-subtracted induction distinguishes cassette-mediated transport (cyanochelins) from siderophore effects shared by both strains.

We observed a slight increase in basal growth of both strains with rising concentration of iron, indicating that addition of iron led to a minor biomass increase. Our background *ΔfutC* mutant was reported^5^ to have low residual ferric iron uptake activity to which we ascribe this observation. In cases of cyanochelin A (see below) and B, there was significant growth of the CctABDEF strain when no FeCl_3_ was added at the highest siderophore concentration (8 μM). We hypothesise that components for media preparation contained residual iron that was subsequently scavenged by the siderophores and the siderophore-accepting strain could utilise it whereas the non-accepting could not.

### The receptor and periplasmic protein are both required for the transport

After successful heterologous expression, we further investigated the minimal set of genes needed to specifically recognise and import cyanochelin B into *Synechocystis* cells. Given the findings above, we hypothesised that the TBDT CctA is not the only component needed for uptake, with the SBP CctB being the main candidate. We therefore constructed seven additional mutant strains bearing various combinations of the genes (see Table 2 and S2).

**Table 2.**
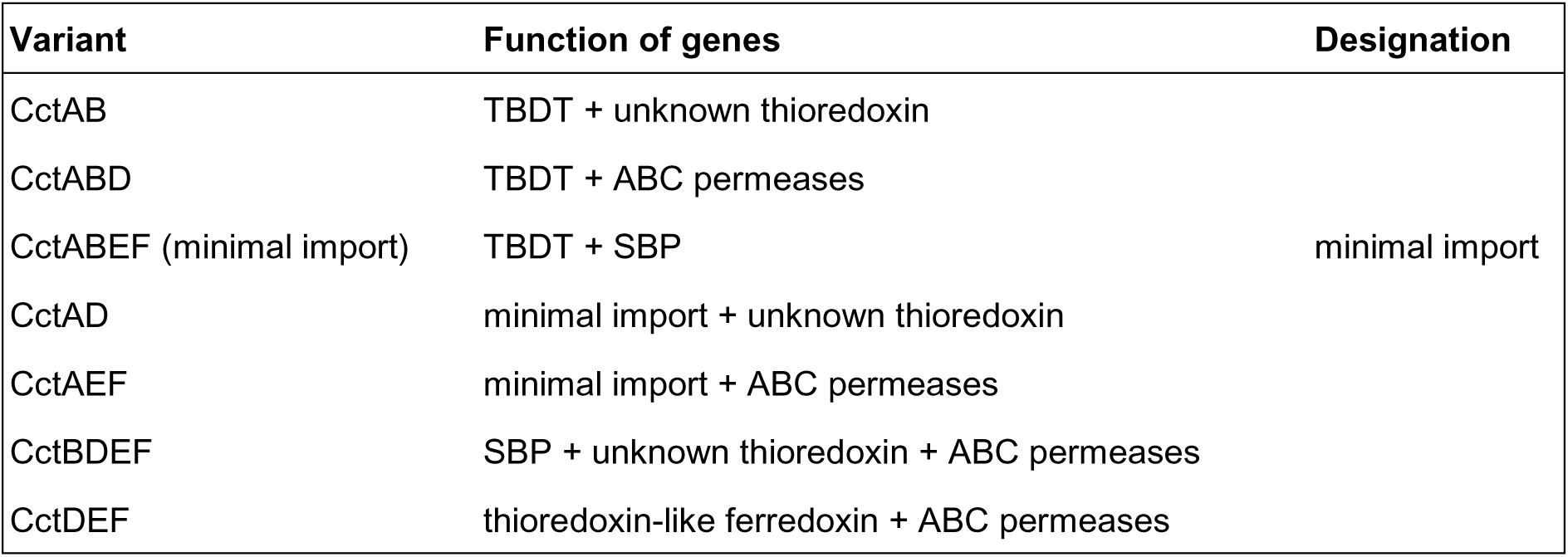
Mutant strains employed in minimal import cassette assay.

In the first set, we combined the CctA with either putative ferredoxin CctD (CctAD) or ABC permeases (CctAEF) or the SBP (CctAB). The second set contained the pair CctA and CctB with either of aforementioned proteins (CctABD, CctABEF). The third was a “negative” set comprising the full import cassette lacking either CctA alone or both CctA and CctB (CctBDEF and CctDEF, respectively). The CctABDEF strain served as a positive control, the *ΔfutC* and the wild-type as negative controls. Iron-starved strains were cultivated under the same conditions as in the initial whole import cassette experiment.

In four variants (CctAD, CctAEF, CctBDEF, CctDEF) the growth was comparable to that of the control *ΔfutC* strain, and these variants were considered negative. Three strains (CctAB, CctABD, CctABEF) outgrew the control and had growth similar to the positive control (CctABDEF), even though the growth extent varied (see Figure 4). We observed growth of strains CctAB, CctABD, CctABEF and positive control in the presence of cyanochelin B even without additional iron as noted above.

**Figure 4.**
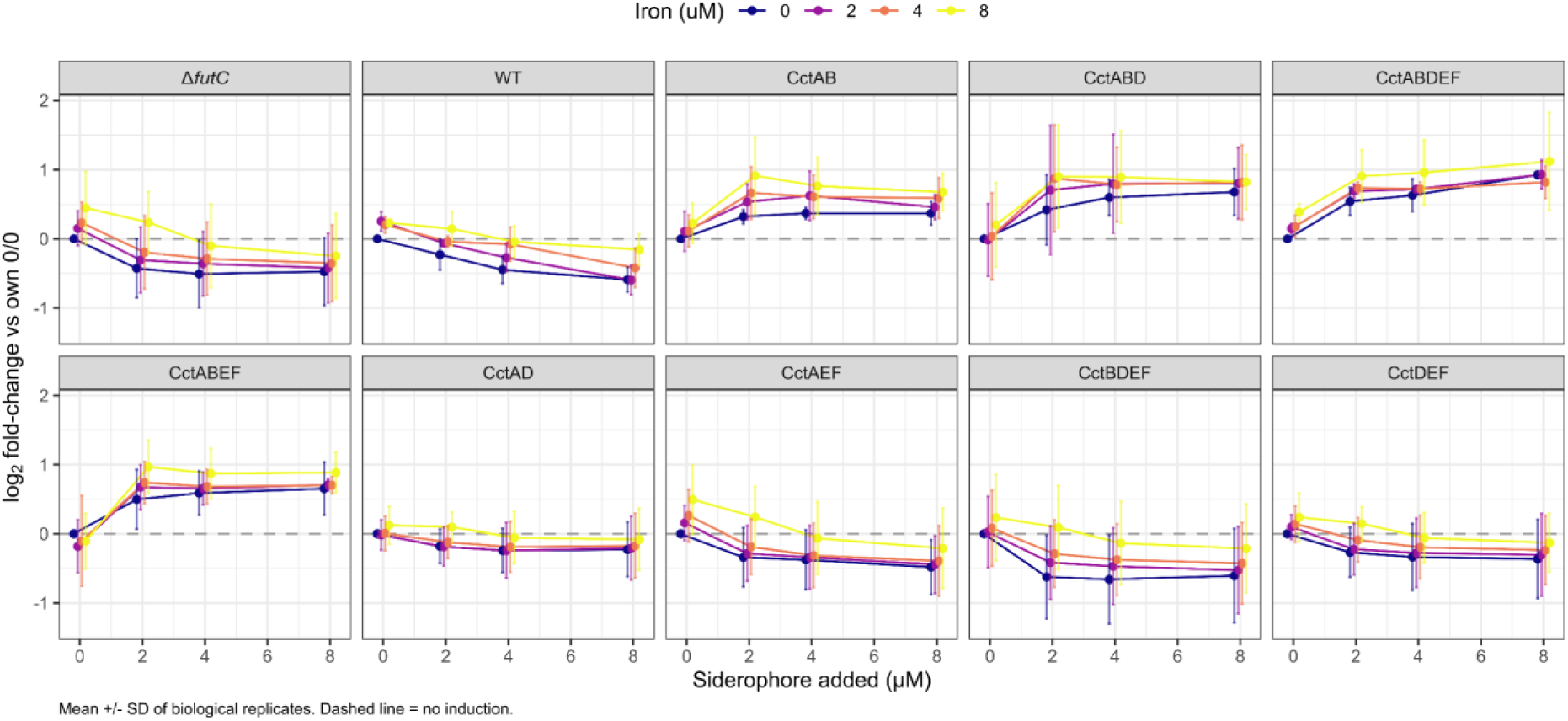
Iron- and siderophore-dependent growth induction in the transporter-variant panel. Growth (OD₇ ₅ ₀ ) is expressed as log₂ fold-change relative to each strain’s own iron- and siderophore-free (0/0) condition, calculated within each biological replicate. Points are the mean of biological replicates (n = 3) and error bars denote ± SD; the dashed line at zero marks no induction. Panels are individual strains; colour encodes the iron concentration (0–8 µM) and the x-axis the added siderophore (0–8 µM). Significance markers are omitted for clarity: statistical comparisons were made by fitting a linear model to the log₂ fold-change values (strain × condition) and contrasting each strain’s induction with that of the ΔfutC control at every iron × siderophore combination using estimated marginal means with Dunnett’s correction; the full contrast estimates and adjusted P values are given in Supplementary Table S4.

The negative results of variants CctBDEF and CctDEF are not surprising, as they lack the cyanochelin B’s TBDT and were not expected to be positive. On the other hand, they also confirm that no *Synechocystis* endogenous TBDT is capable of cyanochelin B transport in cooperation with exogenous SBP. The same conclusion can be derived from growth inhibition of the wild-type strain.

All three positive variants share the presence of both CctA and CctB. Growth was comparable in most cases. The results indicate that both TBDT and SBP confer specificity of cyanochelin B uptake. CctA alone was not able to reconstitute the growth of the mutant strain regardless of the genes added (CctAD, CctAEF), unless the CctB was present as well (CctABD, CctABEF). Moreover, the combination of these two proteins alone was sufficient for growth which suggests that the remaining transport components (ABC permeases, ABC ATPase, theoretically the ferredoxin) can be provided *in trans* by the heterologous host and do not contribute to specificity of uptake.

### The cyanochelin import system shows limited promiscuity across cyanochelins

Further, we probed the substrate strictness of the transport cassette by cultivating the CctABDEF strain in the presence of four xenosiderophores with distinct iron-chelating moieties (see Figure 5): enterobactin (catecholate), pyoverdine mixture (catechol-type chromophore and hydroxamate), and cyanochelins A and C (α-hydroxycarboxylate). Cultivation followed the design of the cyanochelin B experiment – increasing gradient of iron and siderophore.

**Figure 5.**
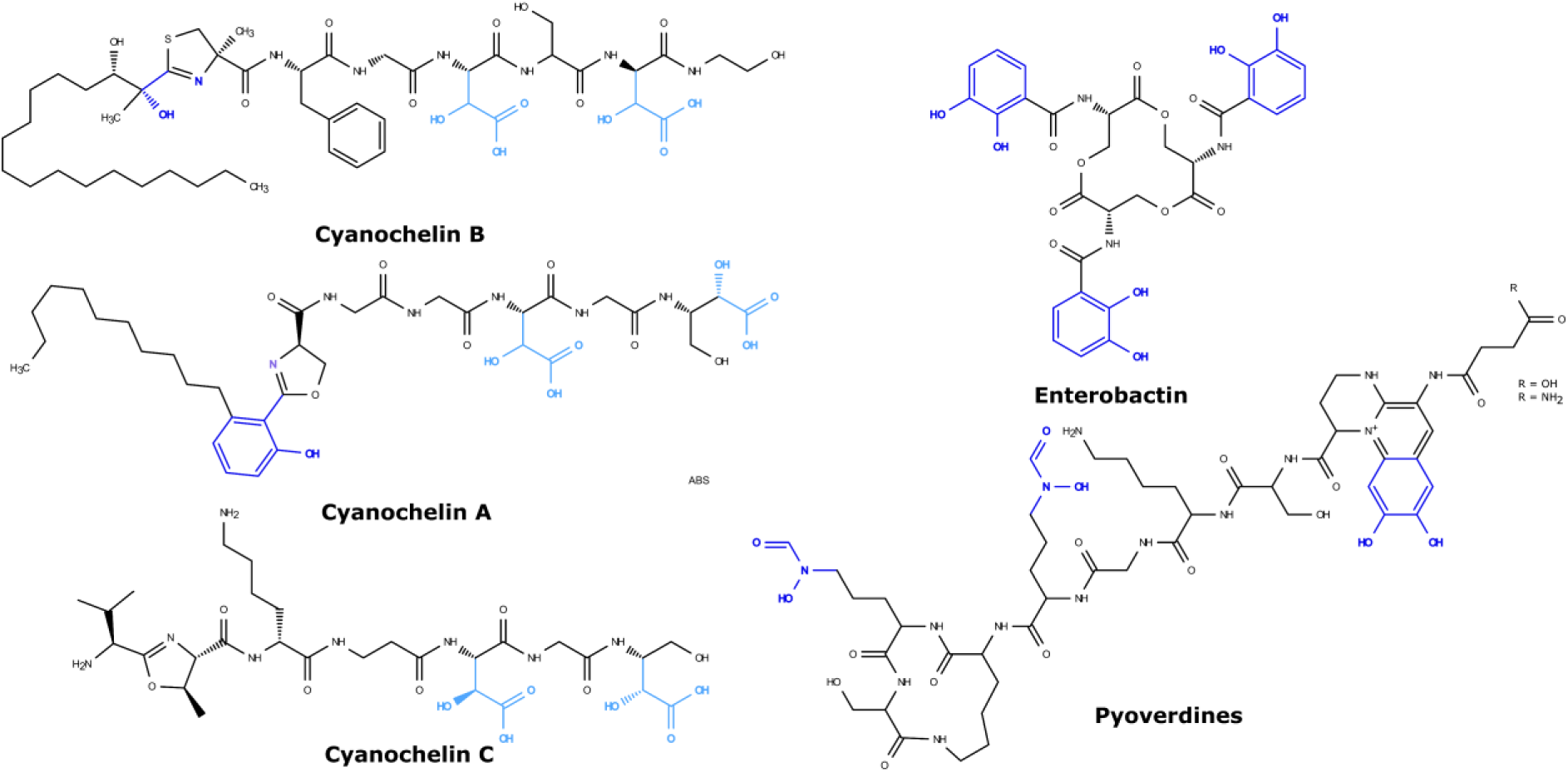
Structures of siderophores used in (xeno)siderophore uptake assay. Light blue colour highlights *β*-hydroxy moieties, dark blue other chelating moieties.

No consistent increase in growth was observed in the presence of cyanochelin C, enterobactin or pyoverdine mixture (see Figure 3C,D,E). Cyanochelin A (Figure 3B), however, was apparently imported by the transport cassette, albeit with lower efficiency: the growth was significantly improved at 8 μM siderophore across all iron levels, and at 4 μM siderophore when combined with 4 and 8 μM FeCl_3_ concentrations. Cyanochelin A, produced by *Rivularia* sp. PCC 7116, was the first cyanochelin to be characterised by our group.^15^ It shares two *β*-hydroxy aspartates for iron chelation with cyanochelin B. Both compounds also possess an aliphatic hydrocarbon chain at the N-terminus of the peptide chain provided by the PKS region of the hybrid PKS-NRPS clusters. The two cyanochelins differ in amino acid sequence: both place a glycine upstream of the first *β*-hydroxy aspartate unit, but the two chelating moieties are separated by serine in cyanochelin B and glycine in cyanochelin A, which also lacks the terminal glycine module that follows the second *β*-hydroxy aspartate in cyanochelin B. Cyanochelin C, produced by *Myxacorys chilensis*, shares no amino acid position outside of *β*-hydroxy aspartate moieties with cyanochelin B and lacks the N-terminal aliphatic moiety. The import cassette therefore discriminates among compounds that share its native chelating group, accepting only the more closely related structure.

### Cyanochelin receptors are phylogenetically related to citrate-hydroxamate siderophore receptors

The specificity of siderophore uptake is predominantly conferred by the outer TBDT and is often restricted to one or few related compounds. Our xenosiderophore assay showed that our studied transport system is capable of transporting structurally similar siderophore cyanochelin A, containing two *β*-hydroxy aspartate groups, but not cyanochelin C that shares this feature as well. Subsequently, we asked what is the evolutionary positioning of the CctA receptor among the cyanobacterial TBDTs.

We retrieved all complete cyanobacterial proteomes from the RefSeq database and used HMM profiles to extract TBDTs. After clustering at 90 % identity, we excluded those that did not possess both plug and barrel domains. As a reference we included several canonical TBDTs such as FhuA or FoxA and also experimentally determined citrate-type hydroxamate transporters for reasons we describe in detail below (see also Table S1). We also included transporters found in the vicinity of cyanochelin-like BGCs identified by Galica *et al*.^15^ This gave us a final dataset of 501 sequences. To capture phylogenetic signal outside strict domain borders,^34^ we used an alignment spanning the N-terminus of the plug domain to the C-terminus of the barrel domain to construct a maximum-likelihood tree (Figure S2). As not all TBDTs include an AMIN domain, we excluded it from the alignment.

CctA is placed within a well-supported Clade E comprising 60 TBDTs (see Figure S2) located in 32 cyanobacterial genomes. Several terminal TBDT branches within this group were obtained, however, the phylogenetic signal did not allow us to reconstruct the mutual relationship between them (Figure 6). We surveyed the respective genomes for the presence of cyanochelin-like BGCs by running antiSMASH. In 14 out of 32 genomes, antiSMASH predicted a NRPS cluster. Eight of these contain a dioxygenase gene or domain, a tailoring enzyme that hydroxylates aspartate to hydroxy aspartate, immediately upstream of the aspartate module, which is the genetic signature of *β*-hydroxy aspartate compounds. After inspection, we found that all eight candidates had already been predicted as cyanochelin-like BGCs by Galica et al. 2021.

**Figure 6.**
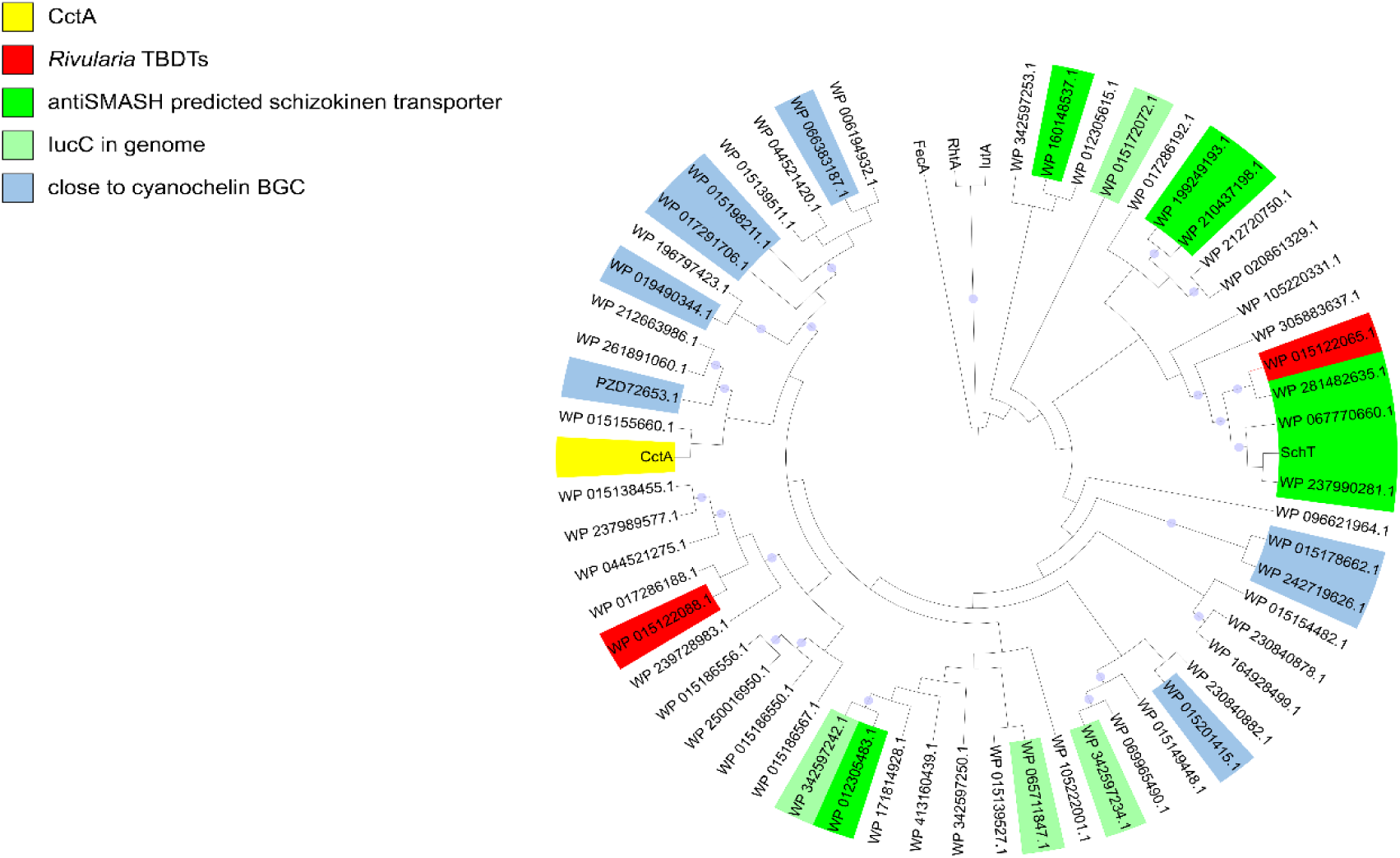
Cladogram of cyanochelin B TBDT branch. Yellow stripe denotes cyanochelin B transporter CctA, blue shows transporters in the vicinity of putative cyanochelin BGCs as characterised by Galica *et al* (2021),^15^ red marks two TBDTs in *Rivularia* sp. PCC 7116 in the vicinity of cyanochelin A BGC that nest inside the branch, bright green denotes putative schizokinen/synechobactin receptors in the vicinity of BGCs, light green TBDTs from proteomes that also include IucC gene as marker for schizokinen production (Table S3). The tree was separately constructed based on Clade E members (see Figure S2) with FecA as an outgroup, branch lengths not shown for clarity. Light blue triangles denote branch support over threshold, SH-aLRT > 80 and UFBoot > 95. Cladogram visualised in iTOL.^36^

Although cyanochelin A transport system has not been yet functionally tested, two out of three TBDTs flanking the cyanochelin A BGC (WP_015122065 and WP_015122088) sit in the same Clade E as CctA. In contrast, the TBDT in the vicinity of the putative cyanochelin C cluster is phylogenetically distant, falling deeper in the reconstructed phylogeny (Figure S2). Phylogenetic proximity thus parallels the experimental result - close for the imported cyanochelin A and distant for rejected cyanochelin C. This may suggest compounds’ relatedness, even though no general conclusion can be drawn.

In the first phylogenetic tree constructed (not shown), five transporters nested in the Clade E were positioned near BGCs likely encoding schizokinen-like compounds in their respective genomes. Schizokinen is a hydroxamate-type siderophore synthesised by type IV non-ribosomal synthesis (often called NRPS-independent siderophore synthetases, NIS).^35^ To further shed a light on the relation of schizokinen and cyanochelin transporters, we constructed another tree (Figure 6 and S2) in which we included the schizokinen receptor SchT, the aerobactin receptor IutA, the rhizobactin 1021 receptor RhtA and the dicitrate transporter FecA. The schizokinen receptor SchT from *Anabaena* sp. PCC 7120 was positioned in the Clade E, being well grouped with other presumed schizokinen transporters, including one of the cyanochelin A potential transporters WP_015122065.1. Whole protein sequence identity between SchT and WP_015122065.1 is 86.1 %, suggesting the latter is schizokinen-like rather than a cyanochelin transporter. Aerobactin and rhizobactin 1021 receptors clearly formed a separate sister branch to the cyanochelin branch, together distinct from the rest of the dataset, while FecA formed a long branch, effectively acting as an outgroup to clades E, F and G. Overall, the results above suggest relatedness between transporters of compounds with entirely different biosynthesis logic, ones being synthesised by the PKS-NRPS pathway, the others with NIS.

To further confirm this finding, we aimed to determine the prevalence of schizokinen-like clusters in cyanobacteria. We screened all proteomes in the phylogenetic tree for the presence of the *Anabaena* sp. PCC 7120 IucC protein (WP_044520601), encoded by one of the essential genes in schizokinen synthesis. All positive hits without exception came from strains that do have a transporter positioned in the CctA clade, further indicating correct nesting of schizokinen type receptors into the clade (see Table S3).

## Discussion

TBDTs as a cornerstone of siderophore import into cells of gramm negative bacteria are of immense importance due to their decisice role in assuring selective transport of the endogenous sideropore or ability to exploit siderophores produced by other members of the community. Therefore understanding of specificity of particular TBDTs has a great value to provide insight into interactions within complex microbial communities.

TBDT-mediated iron uptake in gram negative bacteria and specificity of TBDT transporters is examined via methods such as gene disruption, iron-responsive transcriptomics, labelled iron uptake and others.^37–39^ Despite cyanobacteria being prominent group of photoautotrophic microorganims responsible for large fraction of carbon cycle, the mechanistic studies on siderophore uptake via TBDT are scarse. Recent examples can be found only in a few model genera, such as *Anabaena*,^40^ *Synechocystis*^32^ or *Synechococcus*.^41^ Most non-model cyanobacterial strains remain inaccessible to gene manipulation techniques which limits research on their siderophore transport systems; this is the gap we addressed by employing heterologous reconstitution in *Synechocystis*. *Synechocystis* is promising system because of well-established genetic manipulation methods, lack of an endogenous siderophore production, and presence of only a few TBDTs that could interfere with the analysis. Siderophores known to be utilised by *Synechocystis* are schizokinen and SAV (*Anabaena variabilis* ATCC 29413 siderophore),^42^ desferrioxamine B and aerobactin.^31^ The *ΔfutC* mutant strain cannot rely on the reductive system,^5^ increasing the chance for iron uptake via the supplemented siderophore. Incorporation of five genes putatively involved in the siderophore uptake into the genome of *Synechocystis* sp. *ΔfutC* strain reconstituted growth of an iron-starved culture, established that the cassette does transport cyanochelin B and that *Synechocystis* is capable of heterologously expressing and utilising a xenosiderophore transport system. The extent of its ability remains to be tested with additional transport systems.

A unique cyanobacterial innovation is the fusion of an AMIN domain to the N-terminus of the TBDT. The function of the domain has been studied mainly in two other architectural contexts. The AMIN domain was shown to localise the AmiC amidase to the septal ring during the cell division in bacteria like *E. coli* and *Neisseria gonorrhoeae*. ^26,43^ Moreover, in the cyanobacterium *Nostoc punctiforme* the AmiC2 AMIN domain localises the protein to the maturing septum, where it forms nanopores for cell to cell communication.^44^ The second well-known occurrence is in type IV PilQ secretins in organisms such as *P. aeruginosa* or *Myxococcus xanthus*, where they are also thought to localise monomer units to cell poles during the cell division and possibly anchor the folded transporters to the cell wall.^45,46^ The cyanobacterial cell wall is in many aspects unique among bacteria. Its peptidoglycan layer is thicker than usual for Gram-negative bacteria (10-35 nm (700 nm in extreme) compared to 2-6 nm), with more cross-linking between the chains (56-63 % compared to 20-33 %).^47^ Despite the lack of knowledge, a function similar to that of AmiC and secretins can be hypothesised in AMIN-TBDTs, although its precise role awaits experimental testing. The example of *Nostoc punctiforme* shows that in cyanobacteria, the AMIN domain can localise outer membrane proteins to specific sites in the envelope. Further research is needed to determine whether the AMIN domain interacts with peptidoglycan only, as can be derived from its function in other proteins, or it also serves a purpose unique to cyanobacteria. Notable is also its absence from *Gloeobacterales*. We consider the only representative to be acquired probably via horizontal gene transfer as it is located in a siderophore-receptor-rich genomic island. Therefore, the fusion (or fusions) might have occurred after *Gloeobacterales* diverged from the rest of cyanobacteria.

Our study revealed two essential genes for the transport – TonB-dependent transporter and substrate-binding protein. The TBDT, as the specificity guardian at the outer membrane, is discussed below. Though not exclusively, SBPs are thought to be the main drivers of specificity of transport across the plasma membrane through ligand binding/release kinetics and the accompanying conformation change.^48^ Our observation is consistent with the case of *Synechococcus*, where deletion of a FecB homologue immediately downstream of the probable synechobactin TBDT severely impaired growth of the mutant.^41^ The SBP being essential indicates that the ferri-cyanochelin B is transported into the cytoplasm. In Gram-negative bacteria, the SBP either binds reduced Fe^2+^ in the periplasm after dissociation from the ferri-siderophore complex to prevent precipitation and delivers it to the plasma membrane transporter, or it binds the whole iron-siderophore complex and delivers it to the ABC transporter, with the iron reduction taking place in the cytoplasm.^19^ *Synechocystis*, in reaction to iron depletion, expresses ferrous iron transporter FeoB that was shown to import Fe^2+^ dissociated from siderophores or citrate by reduction in other bacteria.^49–53^ The mutants bearing TBDT without SBP did not grow, which suggests the ferrous iron was not obtained in the periplasm by siderophore hydrolysis, photolysis that would result in Fe^2+^ as was reported for schizokinen.^42^

The ABC permeases and the thioredoxin-like protein were not required in our system. While absence of the permeases can be explained by complementation by the host, the function or necessity of CctD, which is recurrently associated with cyanobacterial siderophore transport loci, remains unresolved. Although the protein’s superfamily is spread through fungi, plants and bacteria (within bacteria, they mostly occur in phylum *Cyanobacteriota*; among fungi mostly in *Ascomycota*), its function has in most cases not been determined experimentally. A few studies have revealed a role in maintaining redox status in yeast cells.^54^

Our assay was designed as a functional reconstitution experiment rather than a direct biochemical measurement of siderophore transport. Therefore, the growth rescue observed upon cyanochelin B supplementation should be interpreted as evidence for cyanochelin B dependent iron acquisition in the *Synechocystis* expression background. Direct quantification of ferri-cyanochelin uptake, intracellular siderophore accumulation or transport kinetics will be required to define the biochemical parameters of the system. Nevertheless, the strict dependence of growth rescue on the presence of both CctA and CctB, together with the negative response to unrelated siderophores, supports the conclusion that these two proteins confer the major substrate-specific steps of the uptake pathway.

Characterised receptors span a spectrum: some TBDTs have relaxed specificity and transport several structurally related compounds, while others are highly strict. For example, *P. aeruginosa*’s enterobactin transporter PfeA has no other known substrate, while PirA transports a wider range of other catechols like dopamine, epinephrine or luteolin as well,^37,55^ and *E. coli*’s FepA additionally transports semisynthetic tris-catecholate myxochelin C.^56^ Pyoverdines are a diverse group of compounds sharing a dihydroquinoline chromophore, but differing in the side chain bound to the chromophore and the peptide chain that varies in length and is specific to a particular strain.^57,58^ The pyoverdines are transported primarily via the FpvA, secondarily via the FpvB receptor.^59^ The peptide chain of the pyoverdines is the specificity-conferring component used for “siderotyping” of the *Pseudomonas* strains into several groups based on pyoverdine structure and its cognate receptor.^60,61^ The FpvAs coevolve with their pyoverdine variants.^62^ The coevolution is driven not only by interspecies, but also intraspecies competition. Different pyoverdine producers synthesise competing pyoverdine forms and many producers and non-producers alike express multiple FpvA variants to utilise the pyoverdines of other strains.^63,64^ Therefore, they accept only their native pyoverdine and/or the closest structural relatives with a similar peptide chain.^65^ On the other hand, FpvB, originally reported as a secondary pyoverdine receptor in *P. aeruginosa*,^59^ was shown to transport at least nine hydroxamate-type siderophores, few of them exclusively, and at least three of those binding to the same cavity.^39,66^ A single TBDT substrate promiscuity is still an under-researched topic, the paradigm being that only close compounds are transported. The example of FpvB opens the question whether there is an evolutionary advantage or process that enables relaxed substrate strictness in TBDTs. The lack of experimental data is currently the main obstacle for better understanding of the phenomenon. In our study, using a phylogenetic approach, we observed that probable transporters of two structurally related siderophores lie in phylogenetic proximity. It could be residual acceptance from a common ancestor. On the other hand, the transporter producer might benefit from slightly relaxed specificity of its transporters, being able to utilise the compounds that diverge moderately, but could outcompete the original in iron pool competition.

Another group of receptors in Clade E were citrate-hydroxamate-type siderophore transporters. *Synechocystis* is known to import schizokinen via a TBDT-ABC pathway. This could partially explain why the whole import cassette did not need to be expressed. The permeases shared higher sequence identity with their homologues in *Leptolyngbya* than the rest of the import genes. However, the schizokinen transporter itself is not able to facilitate utilisation of any cyanochelin.

The specificity of cyanochelin uptake does not revolve only around *β*-hydroxy aspartate iron chelating moieties, as indicated by our results. Cyanochelin A was accepted while cyanochelin C was not. It is worth noting that cyanochelin C *β*-hydroxy aspartate modules do not contain an epimerisation domain. Stereoselectivity has been shown to be an important feature in substrate recognition as demonstrated in pyochelin enantiomers^67^ but remains unresolved for now in cyanochelins. All three cyanochelins also contain a heterocycle at the beginning of the peptide chain. The most notable difference between cyanochelins A and B, accepted by the receptor, and the non-accepted cyanochelin C, is the absence of an acyl chain at the N-terminus of the peptide chain. For the moment, it is not known whether acyl chains influence siderophore recognition. Many amphiphilic lipopeptide siderophores are associated with marine bacteria where the fatty acid is thought to limit diffusive loss of the siderophore in the open ocean.^68,69^ Notable examples, the marinobactins and amphibactins, can have various acyl chain lengths, which can also be cleaved by endogenous hydrolases in the case of marinobactin.^70^ In *Mycobacterium tuberculosis*, carboxymycobactin with a shorter acyl chain is secreted into the extracellular space, while mycobactin with a long acyl chain is attached to the cell wall and transfers iron across the unusual mycobacterial cell envelope.^71,72^ Even though *Leptolyngbya* NIES-3755 is a terrestrial isolate, cyanochelin B’s photolytic properties still favour the anti-diffusion explanation as in the presence of sunlight, the monopolising advantage of siderophore production vanishes.

Our results provide the first insight into the import system of cyanochelins, widely distributed cyanobacterial siderophores. We show that cyanochelin B is transported via the canonical TBDT–ABC route, where in our system the outer membrane receptor and the periplasmic substrate-binding protein are both required for uptake whereas the rest of the pathway could be provided by the host. The TBDT possesses an N-terminal AMIN domain, which is a unique feature of cyanobacterial TBDTs, and is not found in TBDTs of other taxa. We found the import system to be able to transport not only the native substrate, but also the structurally related compound cyanochelin A, whose probable transporter lies in a well-supported phylogenetic branch grouped with schizokinen transporters. We report first expression of a cyanobacterial TBDT in another cyanobacterium, establishing *Synechocystis* as a promising system for xenosiderophore assays.

## Materials and methods

### Culture conditions

All *E. coli* strains were cultivated in LB medium at 37 °C. *Leptolyngbya* sp. NIES-3755 and *Synechocystis* strains were grown in BG-11 medium under ambient temperature and continuous dispersed light.

### Construction of *Synechocystis* strains

Genomic DNA from *Leptolyngbya* NIES-3755 was isolated according to a previously published protocol.^73^ Genes of interest were amplified with Q5 polymerase (New England Biolabs Inc.) using the standard protocol. For primers and plasmids used, see Table S2. PCR clean-up and/or gel purification was done using QIAquick PCR purification Kit or QIAquick Gel Extraction Kit respectively (both QIAGEN). Molecular cloning was performed via NEBuilder HiFi DNA Assembly (New England Biolabs) according to manufacturer’s instructions. For all cloning and plasmid propagation, One Shot TOP10 Electrocomp *E. coli* (Thermo Fisher Scientific) strain was used. Plasmid isolation was performed with QIAprep Spin Miniprep Kit (QIAGEN). Where suitable we changed the start codon to a standard ATG and modified TGA stop codons to TAA codons to better suit *Synechocystis* codon usage.

Parental strain for this study was the *ΔfutC* mutant of *Synechocystis* sp. PCC 6803. The vectors containing genes of interest were introduced into *Synechocystis* via natural transformation. The resulting transformants were segregated by cultivation in increasing erythromycin concentrations. The stability of the insertions was confirmed by PCR.

### Iron uptake assays

Pyoverdine mixture and enterobactin were purchased from Sigma-Aldrich (P8124 and E3910, respectively). Cyanochelins were isolated according to previously described procedures.^15,16^

Iron-starved *Synechocystis* cultures were harvested by centrifugation, washed once with iron-free medium, and resuspended to OD₇ ₅ ₀ = 0.4. Sterile FeCl₃ and siderophore solutions were prepared and diluted to final concentrations of 0, 2, 4, and 8 µM in 96-well plates. Cultures and treatment solutions were mixed in a 1:1 ratio, yielding a starting OD₇ ₅ ₀ of 0.2 in a 200 μl volume. All cultivations were performed in biological and technical triplicates at 25°C under 50 µmol photons m⁻ ² s⁻ ¹ for 10 to 14 days. Growth was assessed by measuring OD₇ ₅ ₀ using a FLUOstar Omega plate reader (BMG Labtech), and data were analysed using custom scripts in R.

### Bioinformatic methods

For the BGCs analysis and prediction, antiSMASH version 8.0.4^74^ was used.

For creation of TBDT datasets, all complete bacterial and cyanobacterial proteomes were retrieved from the NCBI RefSeq database on January 26, 2026, using the NCBI Datasets command-line tool (taxid 2 and 1117, respectively). In both datasets, the TBDT sequences were extracted from each proteome using HMMER (v3.4) searched against the TonB-dependent receptor β-barrel domain profile PF00593, downloaded from Pfam, using a relaxed bit-score threshold of 25 to capture divergent sequences. Duplicate sequence identifiers occurring across multiple genome assemblies were removed using a custom bash script, yielding a final non-redundant set of cyanobacterial TBDT sequences used for all downstream analyses.

For AMIN domain analysis, HMM profiles of the AMIN domain (PF11741) were employed in HMMER (v3.4) to find AMIN-TBDT hits and compared to hits in the cyanobacterial dataset (see below). For distribution in cyanobacteria, the lineages of the taxa bearing the TBDTs were retrieved from NCBI using a custom script.

To construct phylogenetic trees, the cyanobacterial TBDT sequences were clustered at 90% amino acid identity using CD-HIT (v4.8.1). Using HMMER (v3.4), plug and barrel HMM profiles were used to scan through the dataset and domain hits were used to extract plug to barrel region sequences with a custom Python script. Plug and barrel domain sequences from both datasets were independently aligned using MAFFT (v7.525) in L-INS-i mode, and poorly aligned or gappy columns were subsequently removed with trimAl (v1.5.rev0) using automated trimming parameters. Maximum likelihood phylogenetic inference was performed independently for each alignment using IQ-TREE (v3.0.1), with the best-fit substitution model selected from the Q.PFAM matrix set. Branch support was assessed by 1,000 ultrafast bootstrap replicates combined with SH-aLRT tests, and tree topologies were compared between the subsampled and full datasets to evaluate the effect of sequence redundancy on phylogenetic resolution.

## Supporting information

Supplementary tables, each table as a reparate sheet

## Acknowledgement

This work was financed by Czech Science Foundation (22-05478S - Iron monopolization versus community service: the two faces of cyanobacterial beta-hydroxy aspartate lipopeptides) - PI: Pavel Hrouzek. Additional support was provided by the Grant Agency of the University of South Bohemia (GAJU), project no. 126/2024/P, “Do heterotrophic bacteria hijack cyanobacterial siderophores via specific transport mechanisms?” (PI: Berness Peter Falcao). Additional support was provided by OP JAK project “Photomachines” Reg. No CZ.02.01.01/00/22_008/0004624 (Czech Ministry of Education, Youth, and Sports (MEYS)).

## References

1. Keren, N., Aurora, R. & Pakrasi, H. B. Critical Roles of Bacterioferritins in Iron Storage and Proliferation of Cyanobacteria. Plant Physiol. 135, 1666–1673 (2004).

2. Qiu, G.-W., Koedooder, C., Qiu, B.-S., Shaked, Y. & Keren, N. Iron transport in cyanobacteria – from molecules to communities. Trends Microbiol. 30, 229–240 (2022).

3. Andrews, S. C., Robinson, A. K. & Rodríguez-Quiñones, F. Bacterial iron homeostasis. FEMS Microbiol. Rev. 27, 215–237 (2003).

4. Fujii, M., Dang, T. C., Rose, A. L., Omura, T. & Waite, T. D. Effect of Light on Iron Uptake by the Freshwater Cyanobacterium *Microcystis aeruginosa*. Environ. Sci. Technol. 45, 1391–1398 (2011).

5. Katoh, H., Hagino, N., Grossman, A. R. & Ogawa, T. Genes Essential to Iron Transport in the Cyanobacterium *Synechocystis* sp. Strain PCC 6803. J. Bacteriol. 183, 2779–2784 (2001).

6. Kranzler, C. et al. Coordinated transporter activity shapes high-affinity iron acquisition in cyanobacteria. ISME J. 8, 409–417 (2014).

7. Kranzler, C., Lis, H., Shaked, Y. & Keren, N. The role of reduction in iron uptake processes in a unicellular, planktonic cyanobacterium. Environ. Microbiol. 13, 2990–2999 (2011).

8. Hider, R. C. & Kong, X. Chemistry and biology of siderophores. Nat. Prod. Rep. 27, 637 (2010).

9. Kramer, J., Özkaya, Ö. & Kümmerli, R. Bacterial siderophores in community and host interactions. Nat. Rev. Microbiol. 18, 152–163 (2020).

10. Plowman, J. E., Loehr, T. M., Goldman, S. J. & Sanders-Loehr, J. Structure and siderophore activity of ferric schizokinen. J. Inorg. Biochem. 20, 183–197 (1984).

11. Gademann, K. Mechanistic Studies on the Tyrosinase- Catalyzed Formation of the Anachelin Chromophore. ChemBioChem 6, 913–919 (2005).

12. Ito, Y. & Butler, A. Structure of synechobactins, new siderophores of the marine cyanobacterium *Synechococcus* sp. PCC 7002. *Limnol*. Oceanogr. 50, 1918–1923 (2005).

13. Boiteau, R. M. & Repeta, D. J. An extended siderophore suite from Synechococcus sp. PCC 7002 revealed by LC-ICPMS-ESIMS. Metallomics 7, 877–884 (2015).

14. Avalon, N. E. et al. Leptochelins A–C, Cytotoxic Metallophores Produced by Geographically Dispersed *Leptothoe* Strains of Marine Cyanobacteria. J. Am. Chem. Soc. 146, 18626–18638 (2024).

15. Galica, T. et al. Cyanochelins, an Overlooked Class of Widely Distributed Cyanobacterial Siderophores, Discovered by Silent Gene Cluster Awakening. Appl. Environ. Microbiol. 87, e03128–20 (2021).

16. Falcao, B. P. et al. Cyanochelin B: a cyanobacterium-produced siderophore with photolytic properties that negate iron monopolization in UV light. Appl. Environ. Microbiol. 91, e02566–24 (2025).

17. Noinaj, N., Guillier, M., Barnard, T. J. & Buchanan, S. K. TonB-Dependent Transporters: Regulation, Structure, and Function. Annu. Rev. Microbiol. 64, 43–60 (2010).

18. Mirus, O., Strauss, S., Nicolaisen, K., Von Haeseler, A. & Schleiff, E. TonB-dependent transporters and their occurrence in cyanobacteria. BMC Biol. 7, 68 (2009).

19. Schalk, I. J. Bacterial siderophores: diversity, uptake pathways and applications. Nat. Rev. Microbiol. 23, 24–40 (2025).

20. Di Matteo, V. et al. Structural and Stereochemical Elucidation of Cyanochelin C, a Siderophore Associated with Novel Class of Cyanobacterial Acyl Hydrolases. Preprint at 10.64898/2026.07.24.740629 (2026).

21. Obando S., T. A., Babykin, M. M. & Zinchenko, V. V. A Cluster of Five Genes Essential for the Utilization of Dihydroxamate Xenosiderophores in Synechocystis sp. PCC 6803. Curr. Microbiol. 75, 1165–1173 (2018).

22. Drew, D., North, R. A., Nagarathinam, K. & Tanabe, M. Structures and General Transport Mechanisms by the Major Facilitator Superfamily (MFS). Chem. Rev. 121, 5289–5335 (2021).

23. Staron, P., Forchhammer, K. & Maldener, I. Structure–function analysis of the ATP- driven glycolipid efflux pump DevBCA reveals complex organization with TolC/HgdD. FEBS Lett. 588, 395–400 (2014).

24. Hrebicek, O. et al. CluSeek: Bioinformatics Tool to Identify and Analyze Gene Clusters. Preprint at 10.1101/2025.09.16.676505 (2025).

25. De Souza, R. F., Anantharaman, V., De Souza, S. J., Aravind, L. & Gueiros-Filho, F. J. AMIN domains have a predicted role in localization of diverse periplasmic protein complexes. Bioinformatics 24, 2423–2426 (2008).

26. Rocaboy, M. et al. The crystal structure of the cell division amidase AMIC reveals the fold of the AMIN domain, a new peptidoglycan binding domain. Mol. Microbiol. 90, 267–277 (2013).

27. Braun, V., Hartmann, M. D. & Hantke, K. Transcription regulation of iron carrier transport genes by ECF sigma factors through signaling from the cell surface into the cytoplasm. FEMS Microbiol. Rev. 46, fuac010 (2022).

28. Braun, V. Substrate Uptake by TONB - Dependent Outer Membrane Transporters. Mol. Microbiol. 122, 929–947 (2024).

29. Abramson, J. et al. Accurate structure prediction of biomolecular interactions with AlphaFold 3. Nature 630, 493–500 (2024).

30. Strunecký, O., Ivanova, A. P. & Mareš, J. An updated classification of cyanobacterial orders and families based on phylogenomic and polyphasic analysis. J. Phycol. 59, 12–51 (2023).

31. Kranzler, C., Lis, H., Shaked, Y. & Keren, N. The role of reduction in iron uptake processes in a unicellular, planktonic cyanobacterium. Environ. Microbiol. 13, 2990–2999 (2011).

32. Qiu, G.-W. et al. Outer Membrane Iron Uptake Pathways in the Model Cyanobacterium Synechocystis sp. Strain PCC 6803. Appl. Environ. Microbiol. 84, e01512–18 (2018).

33. Bučinská, L. et al. The ribosome-bound protein Pam68 promotes insertion of chlorophyll into the CP47 subunit of Photosystem II. Plant Physiol. pp.00061.2018 (2018) doi:10.1104/pp.18.00061.

34. Gu, S. et al. Feature sequence-based genome mining uncovers the hidden diversity of bacterial siderophore pathways. eLife 13, RP96719 (2024).

35. Dell, M., Dunbar, K. L. & Hertweck, C. Ribosome-independent peptide biosynthesis: the challenge of a unifying nomenclature. Nat. Prod. Rep. 39, 453–459 (2022).

36. Letunic, I. & Bork, P. Interactive Tree of Life (iTOL) v6: recent updates to the phylogenetic tree display and annotation tool. Nucleic Acids Res. 52, W78–W82 (2024).

37. Luscher, A. et al. Plant-Derived Catechols Are Substrates of TonB-Dependent Transporters and Sensitize Pseudomonas aeruginosa to Siderophore-Drug Conjugates. mBio 13, e01498–22 (2022).

38. Will, V. et al. Siderophore specificities of the *Pseudomonas aeruginosa* TonB- dependent transporters ChtA and ActA. FEBS Lett. 597, 2963–2974 (2023).

39. Will, V. et al. The role of FoxA, FiuA, and FpvB in iron acquisition via hydroxamate-type siderophores in Pseudomonas aeruginosa. Sci. Rep. 14, 18795 (2024).

40. Rudolf, M. et al. Multiplicity and specificity of siderophore uptake in the cyanobacterium Anabaena sp. PCC 7120. Plant Mol. Biol. 92, 57–69 (2016).

41. Yong, C.-W., Deng, B., Liu, L.-M., Wang, X.-W. & Jiang, H.-B. Diversity and Evolution of Iron Uptake Pathways in Marine Cyanobacteria from the Perspective of the Coastal Strain *Synechococcus* sp. Strain PCC 7002. Appl. Environ. Microbiol. 89, e01732–22 (2023).

42. Babykin, M. M., Obando, T. S. A. & Zinchenko, V. V. TonB-Dependent Utilization of Dihydroxamate Xenosiderophores in Synechocystis sp. PCC 6803. Curr. Microbiol. 75, 117–123 (2018).

43. Chan, J. M., Hackett, K. T., Woodhams, K. L., Schaub, R. E. & Dillard, J. P. The AmiC/NlpD Pathway Dominates Peptidoglycan Breakdown in Neisseria meningitidis and Affects Cell Separation, NOD1 Agonist Production, and Infection. Infect. Immun. 90, e00485–21 (2022).

44. Büttner, F. M., Faulhaber, K., Forchhammer, K., Maldener, I. & Stehle, T. Enabling cell–cell communication via nanopore formation: structure, function and localization of the unique cell wall amidase AmiC2 of *Nostoc punctiforme*. FEBS J. 283, 1336–1350 (2016).

45. Leighton, T. L., Buensuceso, R. N. C., Howell, P. L. & Burrows, L. L. Biogenesis of *P seudomonas aeruginosa* type IV pili and regulation of their function. Environ. Microbiol. 17, 4148–4163 (2015).

46. Herfurth, M., Pérez-Burgos, M. & Søgaard-Andersen, L. The mechanism for polar localization of the type IVa pilus machine in *Myxococcus xanthus*. mBio 14, e01593–23 (2023).

47. Hoiczyk, E. & Hansel, A. Cyanobacterial Cell Walls: News from an Unusual Prokaryotic Envelope. J. Bacteriol. 182, 1191–1199 (2000).

48. De Boer, M. et al. Conformational and dynamic plasticity in substrate-binding proteins underlies selective transport in ABC importers. eLife 8, e44652 (2019).

49. Katoh, H., Hagino, N., Grossman, A. R. & Ogawa, T. Genes Essential to Iron Transport in the Cyanobacterium *Synechocystis* sp. Strain PCC 6803. J. Bacteriol. 183, 2779–2784 (2001).

50. Kranzler, C. et al. Coordinated transporter activity shapes high-affinity iron acquisition in cyanobacteria. ISME J. 8, 409–417 (2014).

51. Marshall, B., Stintzi, A., Gilmour, C., Meyer, J.-M. & Poole, K. Citrate-mediated iron uptake in Pseudomonas aeruginosa: involvement of the citrate- inducible FecA receptor and the FeoB ferrous iron transporter. Microbiology 155, 305–315 (2009).

52. Ong, A. & O’Brian, M. R. The *Bradyrhizobium japonicum fsrB* gene is essential for utilization of structurally diverse ferric siderophores to fulfill its nutritional iron requirement. Mol. Microbiol. 119, 340–349 (2023).

53. Yeh, T.-Y., Lu, H.-F., Li, L.-H., Lin, Y.-T. & Yang, T.-C. Contribution of fepAsm, fciABC, sbaA, sbaBCDEF, and feoB to ferri-stenobactin acquisition in Stenotrophomonas maltophilia KJ. BMC Microbiol. 25, 91 (2025).

54. Zhang, D. et al. Aim32 is a dual-localized 2Fe-2S mitochondrial protein that functions in redox quality control. J. Biol. Chem. 297, 101135 (2021).

55. Perraud, Q. et al. Opportunistic use of catecholamine neurotransmitters as siderophores to access iron by *PSEUDOMONAS AERUGINOSA* . Environ. Microbiol. 24, 878–893 (2022).

56. Rabsch, W., Voigt, W., Reissbrodt, R., Tsolis, R. M. & Bäumler, A. J. *Salmonella typhimurium* IroN and FepA Proteins Mediate Uptake of Enterobactin but Differ in Their Specificity for Other Siderophores. J. Bacteriol. 181, 3610–3612 (1999).

57. Schalk, I. J., Rigouin, C. & Godet, J. An overview of siderophore biosynthesis among fluorescent Pseudomonads and new insights into their complex cellular organization. Environ. Microbiol. 22, 1447–1466 (2020).

58. Rehm, K., Vollenweider, V., Kümmerli, R. & Bigler, L. A comprehensive method to elucidate pyoverdines produced by fluorescent Pseudomonas spp. by UHPLC-HR-MS/MS. Anal. Bioanal. Chem. 414, 2671–2685 (2022).

59. Ghysels, B. et al. FpvB, an alternative type I ferripyoverdine receptor of Pseudomonas aeruginosa. Microbiology 150, 1671–1680 (2004).

60. De Chial, M. et al. Identification of type II and type III pyoverdine receptors from Pseudomonas aeruginosa. Microbiology 149, 821–831 (2003).

61. Greenwald, J. et al. FpvA bound to non- cognate pyoverdines: molecular basis of siderophore recognition by an iron transporter. Mol. Microbiol. 72, 1246–1259 (2009).

62. Gu, S. et al. Siderophore synthetase-receptor gene coevolution reveals habitat- and pathogen-specific bacterial iron interaction networks. Sci. Adv. 11, eadq5038 (2025).

63. Butaitė, E., Baumgartner, M., Wyder, S. & Kümmerli, R. Siderophore cheating and cheating resistance shape competition for iron in soil and freshwater Pseudomonas communities. Nat. Commun. 8, 414 (2017).

64. Kümmerli, R. Iron acquisition strategies in pseudomonads: mechanisms, ecology, and evolution. BioMetals 36, 777–797 (2023).

65. Hartney, S. L. et al. Ferric-Pyoverdine Recognition by Fpv Outer Membrane Proteins of Pseudomonas protegens Pf-5. J. Bacteriol. 195, 765–776 (2013).

66. Chan, D. C. K. & Burrows, L. L. Pseudomonas aeruginosa FpvB Is a High-Affinity Transporter for Xenosiderophores Ferrichrome and Ferrioxamine B. mBio 14, e03149–22 (2023).

67. Brillet, K. et al. Pyochelin Enantiomers and Their Outer-Membrane Siderophore Transporters in Fluorescent Pseudomonads: Structural Bases for Unique Enantiospecific Recognition. J. Am. Chem. Soc. 133, 16503–16509 (2011).

68. Boiteau, R. M., et al. Siderophore-based microbial adaptations to iron scarcity across the eastern Pacific Ocean. Proc. Natl. Acad. Sci. 113, 14237–14242 (2016).

69. Li, J. et al. Microbial iron limitation in the ocean’s twilight zone. Nature 633, 823–827 (2024).

70. Gauglitz, J. M., Iinishi, A., Ito, Y. & Butler, A. Microbial Tailoring of Acyl Peptidic Siderophores. Biochemistry 53, 2624–2631 (2014).

71. Sritharan, M. Iron Homeostasis in Mycobacterium tuberculosis: Mechanistic Insights into Siderophore-Mediated Iron Uptake. J. Bacteriol. 198, 2399–2409 (2016).

72. Choudhury, M. et al. Iron uptake and transport by the carboxymycobactin-mycobactin siderophore machinery of Mycobacterium tuberculosis is dependent on the iron-regulated protein HupB. BioMetals 34, 511–528 (2021).

73. D’Agostino, P. M., Seel, C. J., Ji, X., Gulder, T. & Gulder, T. A. M. Biosynthesis of cyanobacterin, a paradigm for furanolide core structure assembly. Nat. Chem. Biol. 18, 652–658 (2022).

74. Blin, K. et al. antiSMASH 8.0: extended gene cluster detection capabilities and analyses of chemistry, enzymology, and regulation. Nucleic Acids Res. 53, W32–W38 (2025).

