## Supplementary figures for "Cyanochelin uptake reveals an exclusively cyanobacterial class of AMIN-domain TonB-dependent transporters"

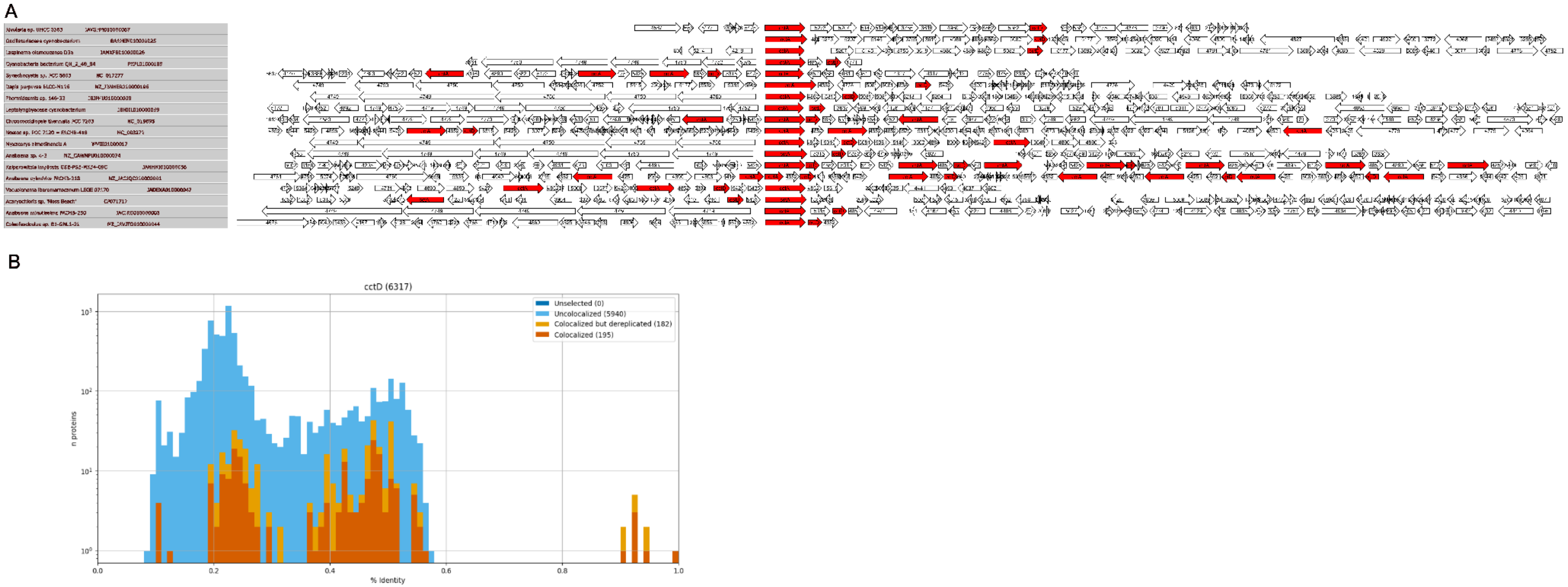


**Figure S1. CluSeek colocalisation analysis of CctA and CctD. A,** Gene composition of CluSeek CctA and CctD colocalisation hits aligned to CctA. Red arrows denote homologues of either CctA or CctD. Both edges were cropped for clarity in Inkscape. **B,** CctD homologue hits in RefSeq. Different bar colours denote CctD homologues uncolocalised with CctA or colocalised and either dereplicated or not. The x-axis shows protein identity (%) of the homologues; the y-axis shows protein count on $\log_{10}$ scale.

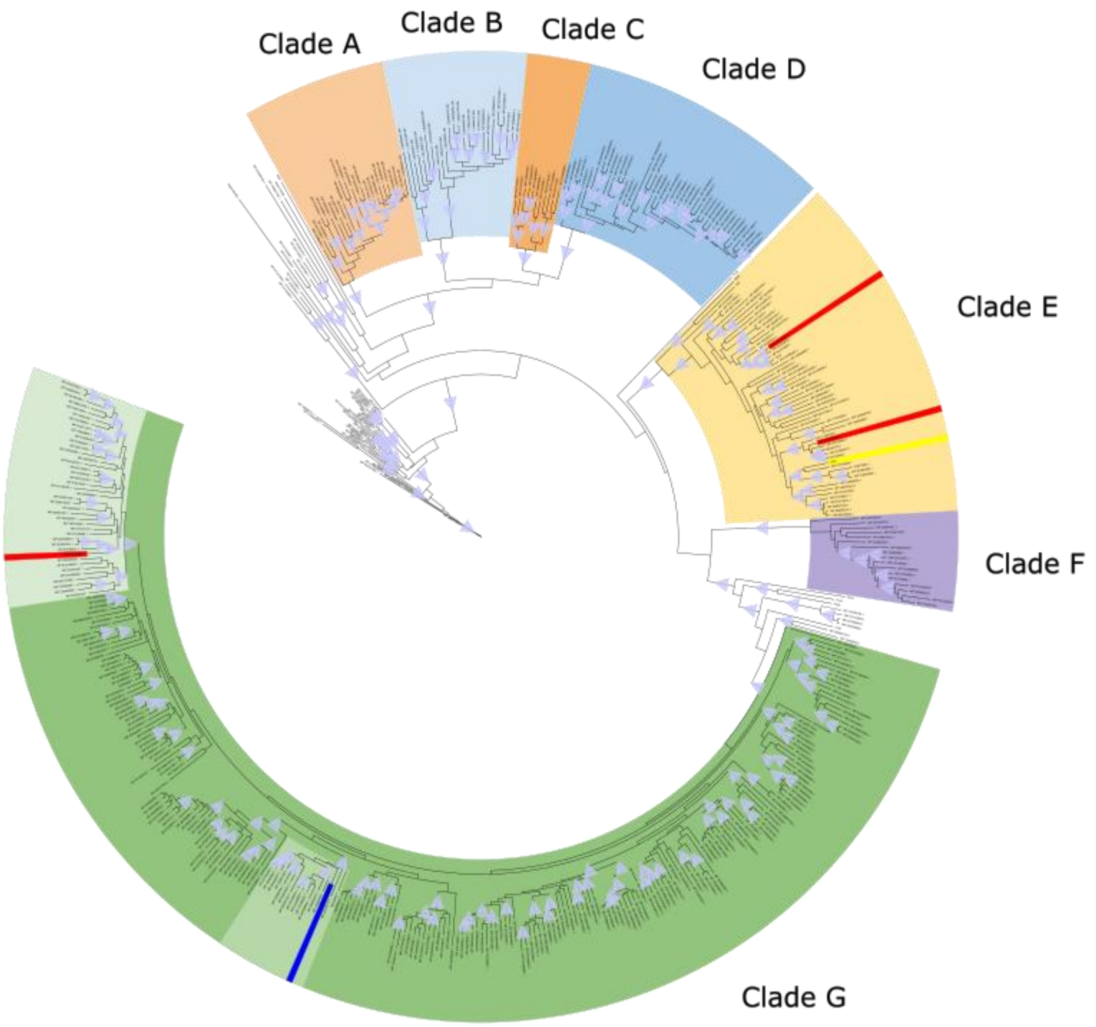


**Figure S2. Phylogenetic tree of cyanobacterial TBDTs.** Extracted sequences were clustered at 0.9 identity. Yellow stripe in Clade E denotes cyanochelin B receptor CctA in *Leptolyngbya*, three red stripes show three TBDTs in the vicinity of cyanochelin A biosynthetic gene cluster in *Rivularia* and dark blue stripe denotes putative cyanochelin C transporter in *Myxacorys chilensis*. Light blue triangles denote branch support over threshold, SH-aLRT > 80 and UFBoot > 95.
